# An RNA-seq quantification method for analysis of transcriptional aberrations

**DOI:** 10.1101/766121

**Authors:** Hiroyuki Kuwahara, Fowzan Alkuraya, Xin Gao

## Abstract

Transcriptome level analysis has been shown to have the great potential for clinical utility. Here, we introduce omega, a between-sample RNA-seq quantification to estimate the abundance level of functional mRNAs which is suitable for analysis of transcriptional aberrations and molecular diagnostics of a range of genetic diseases. By using five diagnosed cases of Mendelian diseases as a case study, we show evidence that omega can improve the signal to detect genes with deleterious transcriptional aberrations and drastically reduce the disease-gene search space.

## Introduction

RNA-seq has been adopted to study a wide range of biological problems. Recent developments have shown the promising use of RNA sequencing (RNA-seq) data in clinical diagnostics (1; 2). Importantly, as transcriptomics helps improve the interpretation of functional consequences of underlying genetic variants, the use of RNA-seq data can lead to the detection of causative genes of genetic disorders that genome-based diagnostics alone has failed to identify (3; 4; 5).

A key step to the application of RNA-seq data is the use of appropriate quantification suitable for specific downstream analysis (6; 7). In transcriptome-based diagnostics, RNA-seq data are typically quantified to detect splicing outliers through analysis of differential splicing events between cases and controls (8; 9). However, a large number of unannotated splicing junctions, often found in RNA-seq data, makes the discovery of causative genes solely from the quantification of splicing junctions challenging. Furthermore, degradation of aberrant transcripts via the activation of the nonsense-mediated decay (NMD) pathway can complicate the prediction of causative genes from splicing outliers. To take into account the degradation of such deleterious variants, expression outliers can be identified via differential expression analysis between cases and controls. Indeed, the integration of these two separate types of transcriptional aberration analysis has shown to increase the diagnostic rate (4; 5). However, because RNA-seq quantifications used for aberrant expression analysis in these previous studies focus on the overall abundance levels of all transcripts (3; 4; 5), they can substantially overestimate the abundance level of functional mRNAs when nonfunctional, aberrant transcripts with splicing error are present, weakening the signal to detect true causative genes.

We hypothesized that transcriptional aberrations can be better evaluated by focusing on the portions of functional mRNAs rather than the overall mRNA abundance levels. That is, we believed that an RNA-seq quantification with the ability to more accurately estimate the abundance level of functional mRNAs is suited for analysis of true expression and splicing outliers. To that end, we have developed omega (observed mRNA’s error-free gene-level abundance), a between-sample RNA-seq quantification that estimates the gene-level abundances of splicing-error-free mRNA transcripts annotated to code for functional proteins. Here, using clinically established cases of genetic disorders, we demonstrate that omega can intensify the signal for deleterious transcriptional aberrations to narrow down the search space for the prediction of causative genes.

## Results and Discussion

### Omega measure

Omega estimates relative abundances of mRNA transcripts produced through error-free splicing processes. Figure 1A overviews a workflow for the omega measure. To compute omega in this workflow, we first estimate the error-free splicing rate and the abundance level of each coding transcript specified in a given transcriptome annotation. We define error-free splicing events to be those without cryptic splicing events based on the annotation in use. We compute the probability of error-free splicing events for each pair of consective exons in each coding transcript based on observed splicing and intron retention events (Fig 1B). For each donor-acceptor pair, we estimate the fraction of error-free mRNA transcripts. And using these values as weights of the count-per-million (CPM) measure of mRNA transcripts for each coding gene, we compute the omega measure (Fig 1C). The decision to use CPM as normalization came from the fact that omega was designed to compare abundances of a given gene between samples (i.e., case and control) and not to compare abundances of different genes within a sample.

**Figure 1.**
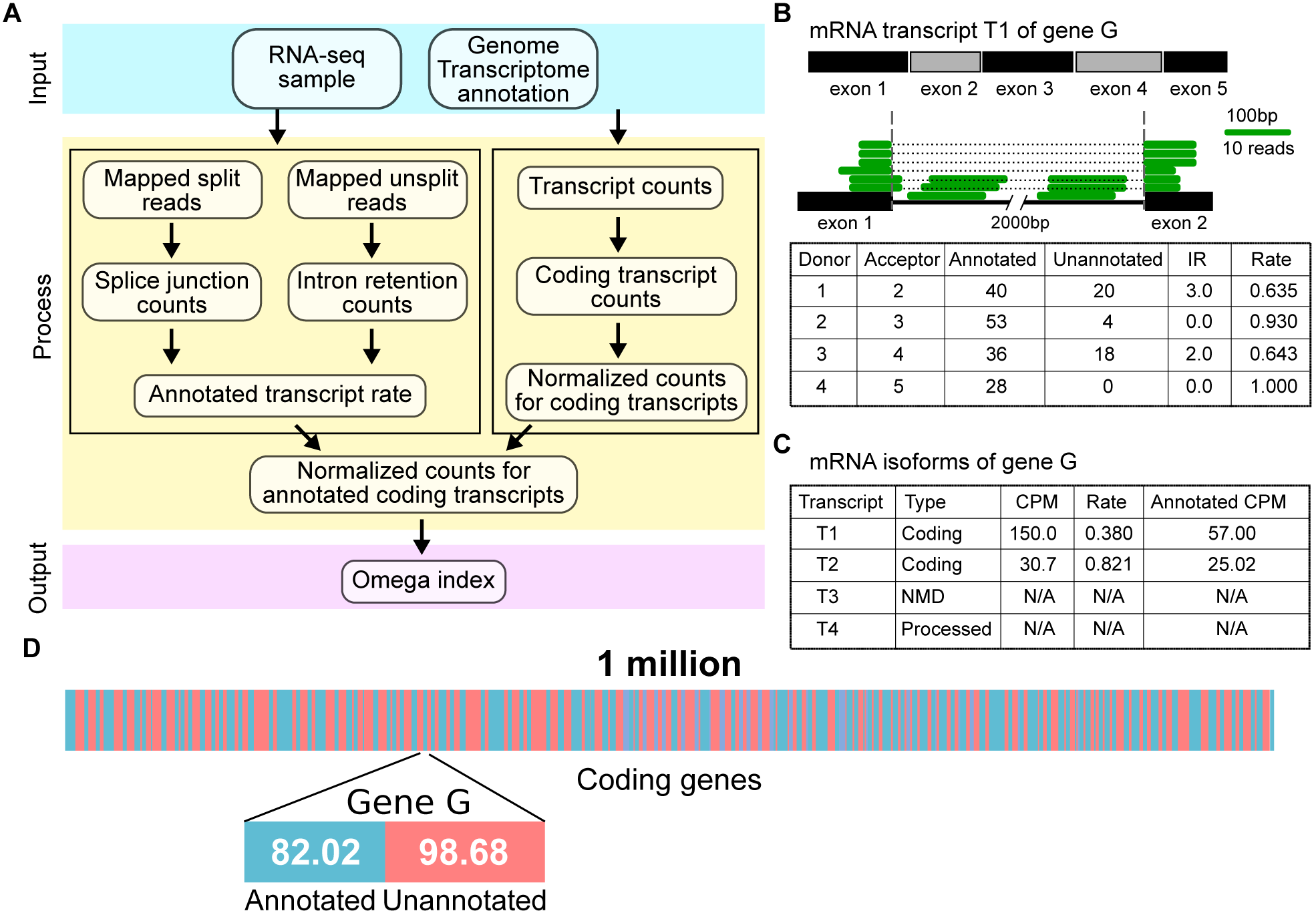
Overview of the omega quantification. (**A**) A workflow to generate the omega measure. (**B**) An illustrative example of the computation of the error-free splicing rate for each donor-acceptor pair in a coding transcript. Here, IR means intron retention. (**C**) An illustration to compute the abundance level of annotated mRNA transcripts for a given coding gene. Here, CPM stands for count-per-million. (**D**) An illustration of the omega measure which partitions the abundance level of each coding gene into those for the annotated, splicing error-free, mRNAs and unannotated, cryptic mRNAs.

The omega measure allows us to partition the expression level of each coding gene into the annotated which represents splicing error-free, functional forms and the unannotated which represents aberrant, nonfunctional forms (Fig 1D). While our definition of error-free splicing depends on a particular annotation that is in use and may not capture every error-free RNA splicing events that constitute functional transcripts, omega can better estimate the relative abundance level of functional mRNAs between samples using the same genome and transcriptome annotation consistently across the board. Assuming that functional mRNAs’ abundances play an important role in controlling the phenotypic state of the cell, the omega measure is expected to differentiate RNA-seq samples based on their phenotypic characteristics. Indeed, clustering analysis of the GTEx data (10) showed that omega outperformed the transcript-per-million (TPM) quantification in differentiating RNA-seq samples with different tissue types (Figure S1).

### Disease-gene prediction

To investigate the value of our new RNA-seq quantification in clinical diagnostics, we applied omega and TPM to five clinically established cases of distinct autosomal recessive disorders with known causal transcriptional aberrations as a case study. RNA-seq datasets of the patients were derived from easily accessible tissue types, skin fibroblasts or lymphocytes. In this study, we used the GTEx RNA-seq data (10) as control and determined low expression outliers out of 19,688 coding genes in the five diagnosed cases using omega and TPM. Here, the desirable outcome from the RNA-seq quantification is the selection of a small set of expression outliers that contains the true disease gene. Because each of these cases has a large number of unique splicing junction sites that are not found in the corresponding cell type of the GTEx cohort (Figure S1), the detection of splicing outliers alone cannot lead to meaningful sets of disease gene candidates, making filtering of candidates given different criteria, such as those based on mRNA abundance levels, essential.

We found that, in all five cases, omega was able to generate smaller sets of outliers that contain the true causal genes than TPM (Fig 2A). For example, in cases 11DG0268 and 11DG0165, while both omega and TPM identified the true causal genes in small sets of outliers, omega was able to filter out non-causal genes further. In addition to the ability to reduce the number of false-positive genes, omega also improved the candidate ranking of true casual genes of these genetic diseases (Table 1). Specifically, COG6, the causative gene of 11DG0268, was ranked 118th in omega and 621st in TPM based on our low expression score (see Methods), while POMT2, the causative gene of 11DG0165, was ranked 13th in omega and 119th in TPM. This indicates that, although some copies of aberrant transcripts of the causative genes were degraded to the level where these genes were considered to be expression outliers, the signal can be further intensified by recognizing other copies of these aberrant transcripts that were escaped from NMD and left in the RNA samples. Indeed, the comparison of the mRNA abundances of the causative genes between omega and TPM highlighted that the consideration of error-free transcripts allowed omega to strengthen the discriminability of the disease-causing gene between the case and the control (Fig 2B). Whereas we showed here the application of omega to autosomal recessive diseases, this quantification is also expected to be suitable for molecular diagnostics of other types of genetic diseases, as well as complex diseases, such as cancer.

**Table 1.** Ranking of the causative genes of the five cases based on the low expression outlier analysis with omega and TPM.

| case ID | causal gene | omega | TPM |
| --- | --- | --- | --- |
| 11DG0268 | COG6 | 118 | 621 |
| 11DG0165 | POMT2 | 13 | 119 |
| 10DG0840 | SPG20 | 2964 | 7173 |
| 16DG1620 | CLCN7 | 495 | 7144 |
| 15DG2154 | DONSON | 429 | 5538 |

**Figure 2.**
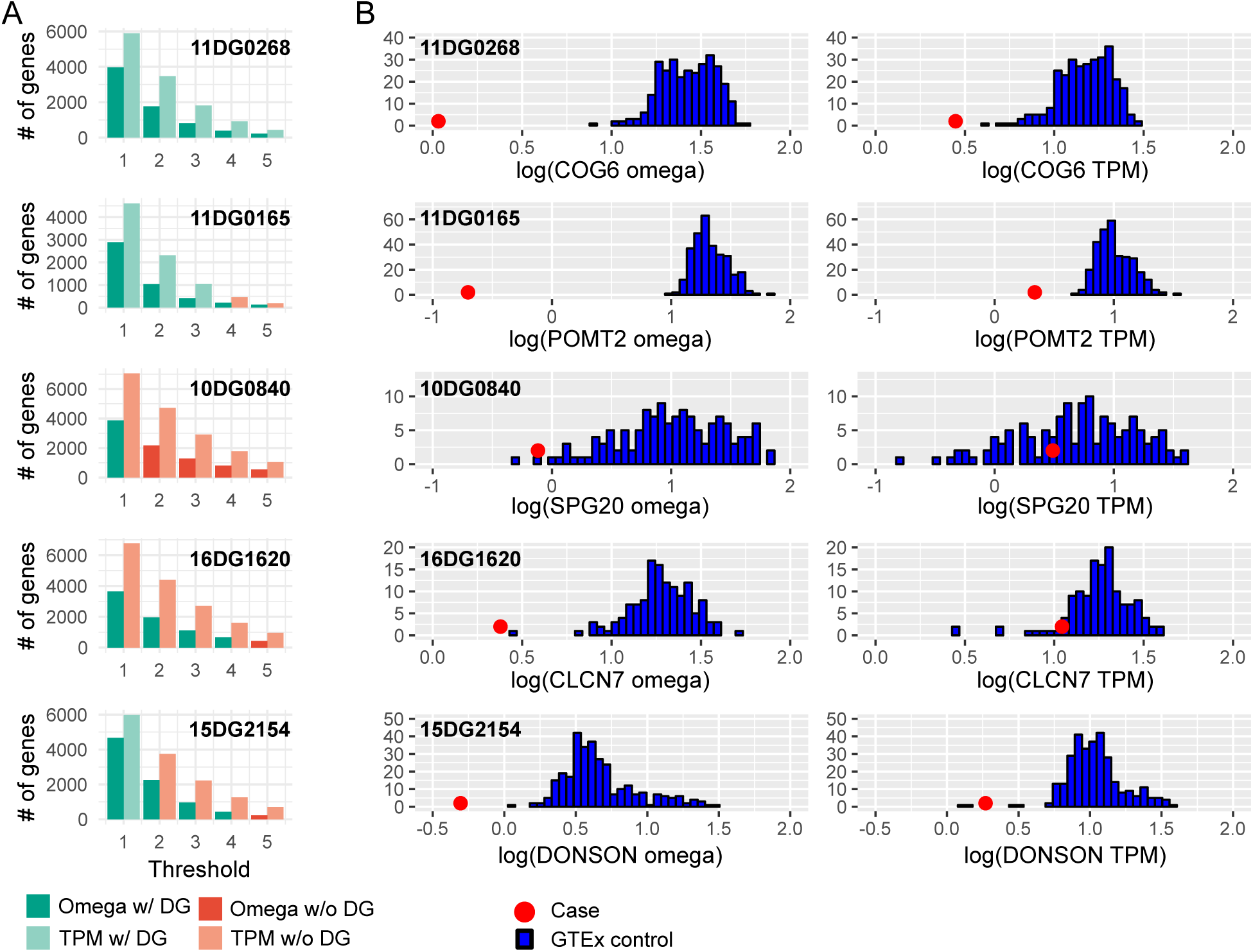
Outlier gene detections in the five cases. (**A**) Histograms showing the number of expression outliers based on various cut-off threshold values with omega and TPM for the five diagnosed cases. See Methods for detail descriptions of the threshold-based selection of expression outliers. Here, DG means disease gene, and each bar is color coded based on the presence and absence of the disease-causing gene in the expression outliers. (**B**) The distribution of the mRNA abundance level of the causal gene for each case using omega and TPM. In each panel, the blue bars indicate a log-transformed abundance-level distribution of the GTEx control, while the red circle represents a log-transformed abundance level from the case.

## Methods

### RNA-seq samples

RNA samples of either fibroblasts or lymphocytes from five patients with clinically established autosomal recessive diseases were prepared meeting all the local ethics guidelines at King Faisal Specialist Hospital and Research Center. We sent these RNA samples to the KAUST core lab for RNA-seq experiments. We determined the quality of each RNA sample based on its RNA Integrity Number (RIN) using Agilent 2100 BioAnalyzer ensured that RNA-seq data were generated from a high-quality RNA sample (RIN ≥ 8.0) for each case. Illumina TruSeq Stranded mRNA was used to prepare the sequencing libraries, and Illumina HiSeq 4000 was used to generate paired-end 150bp reads.

We downloaded GTEx RNASeq samples (10) of the blood and skin tissue types from the Database of Genotypes and Phenotypes with RIN ≥ 6 (n=1,572) and transformed them into the fastq format using SRA Toolkit (https://www.ncbi.nlm.nih.gov/sra/docs/toolkitsoft). These RNA-seq datasets consist of paired-end 75bp reads.

### Error-free splicing probability inference

We aligned the reads to hg38 (GENCODE 25) using STAR 2.6 with the two-pass option (11). We considered reads mapped to chromosomes 1-22 and X in this study. To count the frequency of each splicing junction, we computed the coverage of observed introns from uniquely mapped split reads using SAMtools (12) and BEDTools (13). To classify each observed junction as annotated or unannotated, we computed the distance of each observed splicing junction from the closest donor and acceptor sites using the location of the annotated exons in GENCODE 25. To analyze intron retention events, we used SAMtools and BEDTools to count the number of non-split reads within annotated intronic regions and multiple the counts by the ratio of the read length to the corresponding intron size. Given these measures, we made a Markovian assumption and computed *λ*_*t*_, the probability of error-free splicing for each coding transcript *t*, by

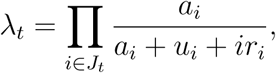

where *J*_*t*_ is a set of splice junctions for transcript *t, a*_*i*_ is the count of the annotated splice junction *i, u*_*i*_ is the count of the unannotated splice junction *i*, and *ir*_*i*_ is the normalized count for the intron retention within the annotated splicing region *i*.

### Transcript count normalization

We used kallisto (14) to map RNA-seq reads to a reference transcriptome for hg38 (GENCODE 25) and to estimate transcript counts. To compute CPM, we selected those transcripts that are annotated as protein-coding from chromosomes 1-22 and X and normalized their counts so that the sum is fixed to 1 million. To compute TPM, we first divided the counts of the selected transcripts by their lengths and normalized the length-normalized counts to have the sum to be 1 million. To obtain the gene-level CPM and TPM for each coding gene, we summed up the normalized value of its coding transcripts.

### Omega

Given the annotated transcript rate and CPM, we computed omega *ω*_*g*_ for gene *g* as follows:

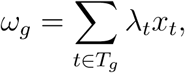

where *T*_*g*_ is a set of coding transcripts annotated for gene *g, λ*_*t*_ is the annotated, splicing-error free, transcription rate for transcript *t* and *x*_*t*_ is the CPM for transcript *t*.

### The definition of low expression outliers

The gene-level mRNA abundance levels of the cases and the GTEx dataset were quantified using TPM and omega, and they were log-transformed with base 10. For each gene *g* in the coding gene set *G*, let *Q*1_*g*_ and *Q*2_*g*_ be the 1st quartile and the 2nd quartile of the log-transformed gene-level mRNA abundance distribution of the GTEx cohort for a given cell type, respectively, and *y*_*g*_ be the log-transformed mRNA abundance level of gene *g* from a case. Then, we computed *a*_*g*_ such that

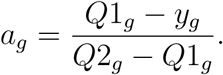

Using a threshold parameter *θ*, we defined low expression outliers to be a subset of *G* in which each gene *g* satisfies *a*_*g*_ ≥ *θ*.

## Acknowledgments

This work was supported by the King Abdullah University of Science and Technology (KAUST) Office of Sponsored Research (OSR) under Awards No. BAS/1/1624-01, URF/1/3412-01, URF/1/3450-01, URF/1/3454-01, FCC/1/1976-18, FCC/1/1976-23, FCC/1/1976-25, FCC/1/1976-26, and FCS/1/4102-02.

## Supplementary figures

**Figure S1.**
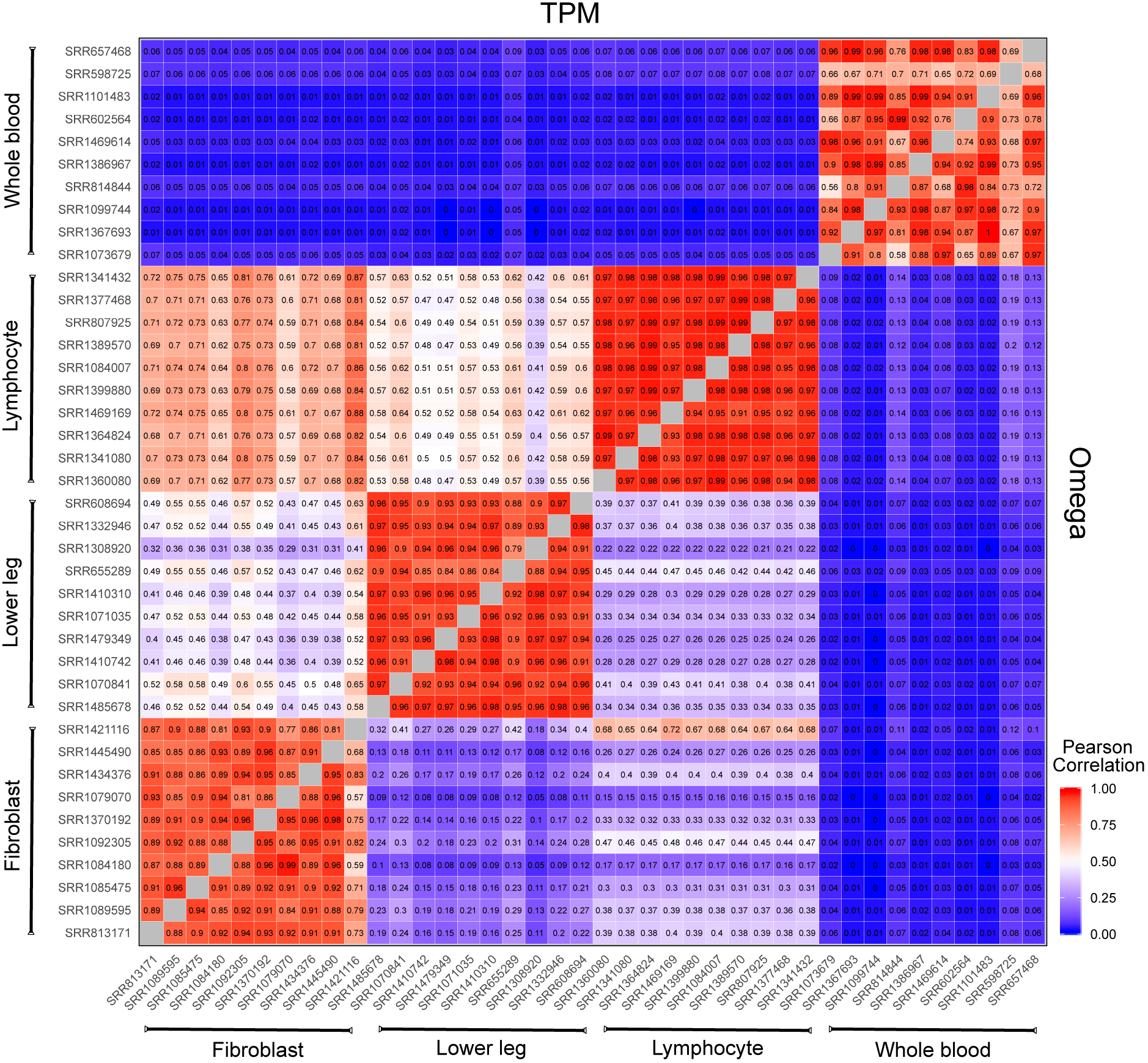
Abundance-based tissue-type clustering using omega and TPM. The upper-left triangular matrix shows the pair-wise correlation with TPM, while the lower-right triangular matrix shows the pair-wise correlation with omega. Ten RNA-seq samples with RIN ≥ 8 were randomly selected from each of 4 tissue types in blood and skin in the GTEx dataset, and pair-wise correlation of these 40 samples was computed using omega and TPM. While both omega and TPM were able to have high correlation values for samples of the same tissue type, omega was able to discriminate samples of different tissue types far better. Specifically, while TPM could not differentiate samples of lymphocytes from those of fibroblasts and lower leg clearly, omega was able to classify those samples based on their tissue types.

**Figure S2.**
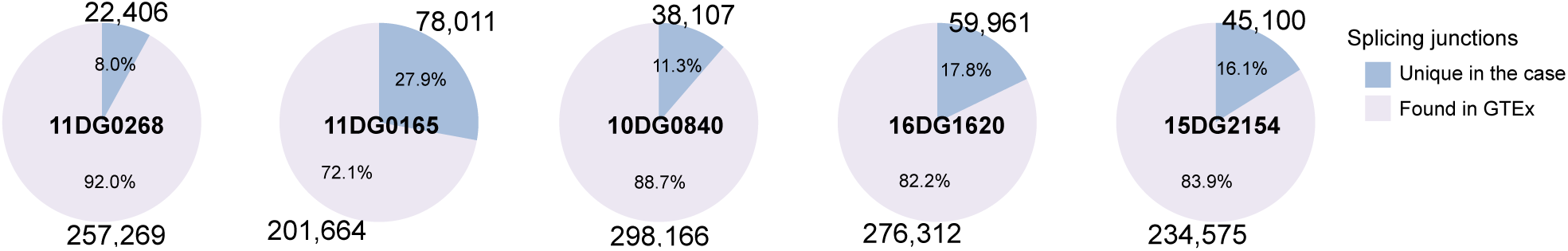
Splicing outliers in the five cases. Splicing junctions found in the RNA-seq samples of the cases were compared against those found in the GTEx samples of the corresponding tissue types (*n* = 133 for lymphocytes and *n* = 305 for skin fibroblasts) to count the number of the junction sites found only in the case and the sites found both in the case and the GTEx control. Splicing junctions in each RNA-seq sample with low read support (*n* < 5) were filtered out.

## References

1. Byron SA, Van Keuren-Jensen KR, Engelthaler DM, Carpten JD, Craig DW (2016) Translating rna sequencing into clinical diagnostics: opportunities and challenges. Nature Reviews Genetics 17: 257–271.

2. Karczewski KJ, Snyder MP (2018) Integrative omics for health and disease. Nature reviews Genetics 19: 299–310.

3. Cummings BB, Marshall JL, Tukiainen T, Lek M, Donkervoort S, et al. (2017) Improving genetic diagnosis in Mendelian disease with transcriptome sequencing. Science translational medicine 9.

4. Kremer LS, Bader DM, Mertes C, Kopajtich R, Pichler G, et al. (2017) Genetic diagnosis of Mendelian disorders via RNA sequencing. Nature communications 8: 15824.

5. Frésard L, Smail C, Ferraro NM, Teran NA, Li X, et al. (2019) Identification of rare-disease genes using blood transcriptome sequencing and large control cohorts. Nature medicine 25: 911–919.

6. Robinson MD, Oshlack A (2010) A scaling normalization method for differential expression analysis of RNA-seq data. Genome biology 11: R25.

7. Evans C, Hardin J, Stoebel DM (2018) Selecting between-sample rna-seq normalization methods from the perspective of their assumptions. Briefings in bioinformatics 19: 776–792.

8. Vaquero-Garcia J, Barrera A, Gazzara MR, González-Vallinas J, Lahens NF, et al. (2016) A new view of transcriptome complexity and regulation through the lens of local splicing variations. eLife 5: e11752.

9. Li YI, Knowles DA, Humphrey J, Barbeira AN, Dickinson SP, et al. (2018) Annotation-free quantification of RNA splicing using LeafCutter. Nature genetics 50: 151–158.

10. Consortium G (2013) The Genotype-Tissue Expression (GTEx) project. Nature genetics 45: 580–585.

11. Dobin A, Davis CA, Schlesinger F, Drenkow J, Zaleski C, et al. (2013) STAR: ultrafast universal RNA-seq aligner. Bioinformatics (Oxford, England) 29: 15–21.

12. Li H, Handsaker B, Wysoker A, Fennell T, Ruan J, et al. (2009) The sequence alignment/map format and SAMtools. Bioinformatics (Oxford, England) 25: 2078–2079.

13. Quinlan AR, Hall IM (2010) BEDTools: a flexible suite of utilities for comparing genomic features. Bioinformatics (Oxford, England) 26: 841–842.

14. Bray NL, Pimentel H, Melsted P, Pachter L (2016) Near-optimal probabilistic RNA-seq quantification. Nature biotechnology 34: 525–527.

